# Compressed Data Structures for Population-Scale Positional Burrows–Wheeler Transforms

**DOI:** 10.1101/2022.09.16.508250

**Authors:** Paola Bonizzoni, Christina Boucher, Davide Cozzi, Travis Gagie, Sana Kashgouli, Dominik Köppl, Massimiliano Rossi

**Affiliations:** University of Milano-Bicocca, Milano, 20126, Italy; University of Florida, Gainesville, 32607, FL, USA; Dalhousie University, Halifax, B3H 4R2, Canada; Tokyo M&D University, Toyko, Japan

**Keywords:** Succinct data structures, Burrows–Wheeler transform, PBWT, Pattern matching

## Abstract

The positional Burrows–Wheeler Transform (PBWT) was presented in 2014 by Durbin as a means to find all maximal haplotype matches in *h* sequences containing *w* variation sites in 𝒪(*hw*)-time. This time complexity of finding maximal haplotype matches using the PBWT is a significant improvement over the naïve pattern-matching algorithm that requires 𝒪(*h*^2^*w*)-time. Compared to the more famous Burrows-Wheeler Transform (BWT), however, a relatively little amount of attention has been paid to the PBWT. This has resulted in less space-efficient data structures for building and storing the PBWT. Given the increasing size of available haplotype datasets, and the applicability of the PBWT to pangenomics, the time is ripe for identifying efficient data structures that can be constructed for large datasets. Here, we present a comprehensive study of the memory footprint of data structures supporting maximal haplotype matching in conjunction with the PBWT. In particular, we present several data structure components that act as building blocks for constructing six different data structures that store the PBWT in a manner that supports efficiently finding the maximal haplotype matches. We estimate the memory usage of the data structures by bounding the space usage with respect to the input size. In light of this experimental analysis, we implement the solutions that are deemed to be superior with respect to the memory usage and show the performance on haplotype datasets taken from the 1000 Genomes Project data.

## 1 Introduction

Haplotype-resolved pangenomes provide a global representation of common variants and genetic variability in whole-genome sequencing of hundreds of thousands of individuals such as the ones collected by the UK Biobank (UKB) [15] or TOPMed projects [31]. Improved haplotype phasing in large cohorts is facilitating the comprehensive study of variations at chromosome-level for genome evolution and clinical applications. In the framework of computational pangenomics, the positional BWT (PBWT), which is a method of permuting the elements of each column of a *h*×*w* binary matrix M[1..*h*][1..*w*], is a key instrument in the compact representation of large haplotypes data sets [9]. Indeed, due to the intrinsic capability of the PBWT of saving space in memorizing haplotype data and even in analyzing large haplotypes panels, it is becoming a relevant data structure for pangenomics (see the recent tutorial on data structures [2]). It is used in relevant computational steps related to haplotype phasing and analysis, such as the matching procedure in reference panels of haplotypes. Moreover, the notion of PBWT has been extended to graph pangenomes representations of haplotypes with the name of Graph positional BWT or GBWT, and it is currently the building block of sequence to graph aligners in the VG toolkit [2].

We note that the original intention of the PBWT was to develop a means to find maximal haplotype matches in a set of *h* sequences, each containing *w* variation sites. In this context, it is assumed that each variation site is bi-allelic, meaning that there exists only two observed alleles at a locus in the genome and no insertions or deletions. Although this binary encoding of genetic information appears to remove a lot of information, it is very common in the analysis of variations of diploid species, where variations are filtered to only contain bi-allelic sites [32, 23]. The PBWT is easy to define and construct. The first column of the PBWT of M is the same as the first column of M. To build column *j* of the PBWT of M, we stably sort the rows of M[1..*h*][1..*j*] in co-lexicographic order (i.e., sorted order from right-to-left) starting at column *j*−1. Thus, the PBWT is something like a radix sort or, for those who know it, like a wavelet matrix [7] on its side. Given the PBWT of M, a pattern *P* [1..*m*], and a starting column *𝓁*, the PBWT allows us to efficiently find all the rows of M which contain *P* between columns *𝓁* and *𝓁* + *m*−1. If there are no such rows, the data structure returns all the rows of M that contain *P* [1..*m*′] between columns *𝓁* and *𝓁* + *m*′−1, where *P* [1..*m*′] is the longest prefix of *P* that occurs in M. Durbin [9] showed that maximal haplotype matches can be found in *O*(*hw*)-time since it amounts to finding set-maximal exact matches (SMEMs) using the PBWT, where a SMEM is defined to be an exact match that is flanked by the right and the left by mismatches between the pattern and the string.

Since its initial development, the PBWT has been applied and extended in numerous ways. It has been used for genotype imputation [26], and to create a genotype database search method that is privacy-preserving (PBWT-sec) [28]. Novak et al. [22] and Sirén et al. [29] used the PBWT to encode a graph for haplotype matching (g-PBWT) and graph pangenome indexing [2]. Sanaullah et al. [27] replaced all arrays with linked lists to define a dynamic version of the PBWT (d-PBWT). The original PBWT has been used to compute all-pairs Hamming distances [20] and for finding all maximal perfect haplotype blocks in linear time [1].

In this paper, we consider the problem of Durbin [9] that aims to find SMEMs in haplotype data using the PBWT, and break this problem into sub-problems that can be efficiently solved using various data structures. In particular, we define eight different smaller data structures that have different functionality, and can be combined to build data structures for storing the PBWT in a manner that SMEMs can be found efficiently. Some of these data structures—such as that of Rossi et al. [25]—exist in the literature in another context, while others are novel. Together, this provides a comprehensive list of data structures for storing the PBWT. We study the theoretical bounds of each component data structure, and benchmark their memory consumption under a practical setting. This benchmarking gives an estimate on how much space our six solutions would take in practice and allows us to conclude which is most likely to be efficient in practice. This conclusion leads us to an implementation of a fully functional data structure that efficiently finds SMEMs in the PBWT. Lastly, we show the utility of our approach by building our data structure and finding SMEMs in increasingly larger datasets from the 1000 Genomes Project data.

Hence, contributions of this work are threefold: (1) a formal definition of finding SMEMs in the PBWT, and the queries needed to support finding SMEMs; (2) a comprehensive study of component data structures used to support the aforementioned queries, and the ways that they can be combined to create data structures to find SMEMs in the PBWT; and lastly, (3) an experimental evaluation of the components and data structures on both simulated data, and the 1000 Genomes Project data. In our experimental evaluation, we first benchmark the memory usage of the various data structures on simulated data to determine those that are going to be scalable to large datasets. Second, we present a full evaluation of the running time and memory usage of those data structures on the 1000 Genomes Project data. Inclusive of this later evaluation, is the evaluation of the time required to find SMEMs in the PBWT.

## 2 Definitions

### 2.1 Basic Definitions

We define a string *S* over a finite, ordered alphabet Σ = {*c*_1_, …, *c*_*ς*_}of *σ* characters to be a sequence of *n* characters *S* = *S*[1..*n*]. We denote the empty string as *ε*. We denote the length of *S* by *S*. We denote the *i*-th prefix of *S* as *S*[1..*i*], the *i*-th suffix as *S*[*i*..*n*], and the substring spanning position *i* through *j* as *S*[*i*..*j*], with *S*[*i*..*j*] = *ε* if *i* > *j*. Given two strings *S* and *T*, we say that *S* is lexicographically smaller than *T* if either *S* is a proper prefix of *T* or there exists an index *i* ≥ 1 such that *S*[1..*i*] = *T* [1..*i*] and *S*[*i* + 1] occurs before *T* [*i* + 1] in Σ. We denote this as *S* ≺ *T*.
We denote a string *S* as *binary* if *σ* = 2. For the sake of simplicity, we assume that all input strings are binary strings defined over Σ = {0, 1} unless stated otherwise. The following definition will be used later in this work. Given a binary character b ∈ Σ, we denote the negation of b as ¬ b = 1 − b.

We denote a matrix of *h* rows and *w* columns as M[1..*h*][1..*w*]. We let col(M)_*j*_ denote the *j*-th column of M, i.e., the string drawn as col(M)_*j*_ = M[1..*h*][*j*] = M[1][*j*]M[2][*j*] *…* M[*h*][*j*].

### 2.2 RMQ, PSV, and NSV

Given an array A[1..*n*] of integers, a range minimum query (RMQ) for two positions *i* ≤ *j* asks for the position *k* of the minimum in A[*i*..*j*], i.e., *k* = argmin_*k*′_ ∈[*i*..*j*]*A*[*k*′]. We denote this query by RMQ_A_(*i, j*). Given a position *i* in *A*, we define the previous-smaller-value (PSV) as PSVA(*i*) = max({*j* : *j* < *i*, A[*j*] < A[*i*]} *∪* {0}). We define the next-smaller-value (NSV) as NSVA(*i*) = min({*j* : *j* > *i*, A[*j*] < A[*i*]} *∪* {*n* + 1}).

### 2.3 Rank and Select

For a string *S*, a character c ∈ Σ, and an integer *j*, the rank query *S*.rank_c_(*j*) counts the occurrences of c in *S*[1..*j*], and the select query *S*.select_c_(*j*) gives the position of the *j*-th c in *S*.

### 2.4 Suffix Array and Burrows–Wheeler Transform

Given an input string *S*, we assume *S* is terminated by a special symbol $ that is lexicographically smaller than any other character in Σ. Next, we consider all rotations of *S* in lexicographical order. We denote the matrix of these rotations, which is commonly referred to as the *Burrows–Wheeler Transform Matrix*, as ℳ. It is convenient to additionally store the starting position (in *S*) of each of these rotations. Since ℳ is stored lexicographically all occurrences of any pattern *P* occur together in a contiguous range, implying finding all occurrences of *P* corresponds to finding the contiguous range in ℳ that begins with *P*. Obviously, ℳ requires 𝒪(*n*^2^)-space but it follows that finding this contiguous range can be accomplished by only storing and using the first column of ℳ (denoted as F) and the last column of ℳ (denoted as L).

The characteristic of ℳ that allows only L and F to be used to find all occurrences of a pattern *P* is the LF-mapping which states that the *i*-th occurrence of a character c in L and the *i*-th occurrence of c in F correspond to the same occurrence of c in *S*. Here, we note that L is referred to as the *Burrows–Wheeler Transform array* and is denoted as BWT. Using the LF-mapping, *S* can be constructed from BWT in linear-time in the length of *n*, and therefore, is deemed to be reversible. The starting positions of all the rotations is a permutation of {1, …, *n*}, and is referred to as the *suffix array* of *S* since it is a lexicographical sort of all the suffixes of *S*. We denote this as SA_*S*_, or just SA if *S* is unambiguously defined. We can define the *Burrows–Wheeler transform (BWT)* with respect to SA as a permutation of the symbols of *S* such that BWT_*S*_ [*i*] = *S*[SA_*S*_ [*i*] −1] if SA_*S*_ [*i*] > 1; otherwise it is $. Again, we denote BWT_*S*_ as simply BWT if *S* can be inferred unambiguously.

### 2.5 LF and FL Mapping via rank and select

If we know the location of a character *S*[*i*] in BWT, we can compute *S*[*i* + 1] using rank and select. More formally, to go from *S*[*i*] to *S*[*i* + 1], we can *forward step* in the BWT, using FL-mapping:

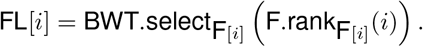

Similarly, to move backward (i.e., from *S*[*i*] to *S*[*i −* 1]) in the BWT using LF-mapping as follows:

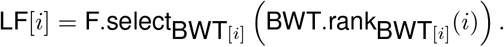

Both LF-mapping and FL-mapping can be efficiently supported using various data structures, including the recent data structures of Nishimoto and Tabei [21].

### 2.6 Longest Common Prefix, Longest Common Suffix, and Longest Common Extension

Given two strings *S* and *T*, we denote the length of the longest common prefix between *S* and *T* as lcp(*S, T*). Using this notation, we define the longest common prefix array of an input string *S* of length *n* as LCP[1..*n*] where LCP[*i*] = lcp(*S*[SA_*S*_ [*i*]..*n*], *S*[SA_*S*_ [*i* − 1]..*n*]) for all *i* > 1, and LCP[1] = 0.

Similarly, given two strings *S* and *T*, we denote the length of the longest common suffix between *S* and *T* as lcs(*S, T*).

Given an input string *S* of length *n* and two integers 1 ≤ *i* ≤ *n* and 1 ≤ *j* ≤ *n*, the Longest Common Extension (LCE) is the longest substring of *S* that starts at both *i* and *j*. Moreover, as we will discuss in this work, there are multiple data structures that efficiently support LCE queries for a string *S*.

### 2.7 Definition of the r-index

Considering matrix ℳ, we see that F can be stored as an integer array of size Σ that stores the number of occurrences of each character in Σ in *S*. Similarly, L (or BWT) can be stored compactly by considering the *runs* of each character. We continue with the convention of denoting the number of runs as *r*. This was first noted by Mäkinen and Navarro [19] in 2005 when they defined the *Run-Length Burrows–Wheeler Transform* (RLBWT), which is a compact representation of the BWT that scales in size with respect to the number of (maximal equal character) runs in the BWT. In the simplest format the RLBWT is an array of that stores pairs storing the character c of the run and the length of the run. Thus, Mäkinen and Navarro demonstrated that the BWT can be stored in 𝒪 (*r*) words while still efficiently supporting queries of the form: find all occurrences of the longest suffix match of a pattern *P* in *S*. However, efficiently storing the SA in a manner that allows all occurrences of these matches in *S* to be found is proved to be more challenging. In order to store the SA efficiently a subset of the SA, i.e., a “SA sample”, is stored and the missing SA values are computed rather than stored. Mäkinen and Navarro showed that the size of SA sample is inversely proportional to the time required to support finding all the occurrences of a pattern *P*, i.e., using a small SA sample results in less-efficient time to find the occurrences of a pattern, and a larger SA sample results in a faster method to find the occurrences of a pattern. Determining how to store the SA in 𝒪 (*r*) words and in a manner that would still efficiently support finding the occurrences was left as an open problem for over a decade.

In 2018 Gagie et al. [13] showed how SA can be sampled at all *r* runs and be combined with a small auxiliary data structure (called Φ) to provide efficient access to SA. Their resulting data structure, called the *r-index*, has been shown to have scalable construction on large repetitive datasets [25]. Hence, the *r*-index consists of the RLBWT, a SA sample at either the beginning or end of each run in the BWT, and Φ that allows recursive access to the missing SA samples; together Gagie et al. demonstrate that not only the number of occurrences of a pattern *P* can be found efficiently but also the location (in *S*) of each of these occurrences.

### 2.8 SMEMs, Matching Statistics, and Thresholds

Given an input string *S* and a pattern *P*, we refer to a set-maximal exact match (SMEM) as an exact match between *S* and *P* that cannot be extended on the left or the right. Bannai et al. [3] demonstrated that SMEMs can be found in the *r*-index augmented with random access to *S*, via the calculation of *matching statistics* for the pattern *P*, which are defined as follows: the matching statistics are an array *A*[1.. |*P*|] of (pos, len) pairs such that (1) *S*[*A*[*i*].pos..*A*[*i*].pos + *A*[*i*].len − 1] = *P* [*i*..*i* + *A*[*i*].len 1]; and (2) *P* [*i*..*i* + *A*[*i*].len] does not occur in *S*. Informally, for each position *i* in the pattern *P*, the *i*-th entry of the array A stores a position in *S* (i.e., pos) where the pattern *P* [*i*.. |*P*|] and *S*[pos.. |*S*|] share a longest common prefix of maximal length (len). Next, we can find SMEMs of *P* with respect to *S* in a left to right pass of the matching statistics. The same authors went on to describe how to compute the matching statistics by using an auxiliary data structure (called *thresholds*) along with the traditional *r*-index and the random-access structure, which indicate a position in ℳ of one of the minimum LCP values1 between consecutive runs of the same character c. The thresholds data structure requires storing 𝒪 (*r*) words—in fact, we can be more precise and state that at most 4*r* copies of 1 are needed to mark the thresholds since we need a bit vector for each character (i.e., A, C, G, and T). This is illustrated in Table 1. Hence, the complete thresholds data structure requires the addition of 𝒪 (*r*)-space, and the *r*-index with thresholds requires 𝒪 (*r*) words of space. Bannai et al. left it open as how to construct the thresholds efficiently. This was resolved in 2022 by Rossi et al. [25] when they demonstrated that this can efficiently constructed with the SA samples and RLBWT via prefix-free parsing. As described by Rossi et al., the *r*-index is currently being explored for aligning reads to a reference genome, which requires finding approximate alignments between exact matches. To find these approximate matches, the sequence in the reference is needed, which is obtained via a data structure that provides random access to the reference, such as a Tabix index [18] or an RLZ parse [17]. Figure 1 illustrates the main components of the *r*-index, i.e., the SA sample and RLBWT.

**Table 1.**
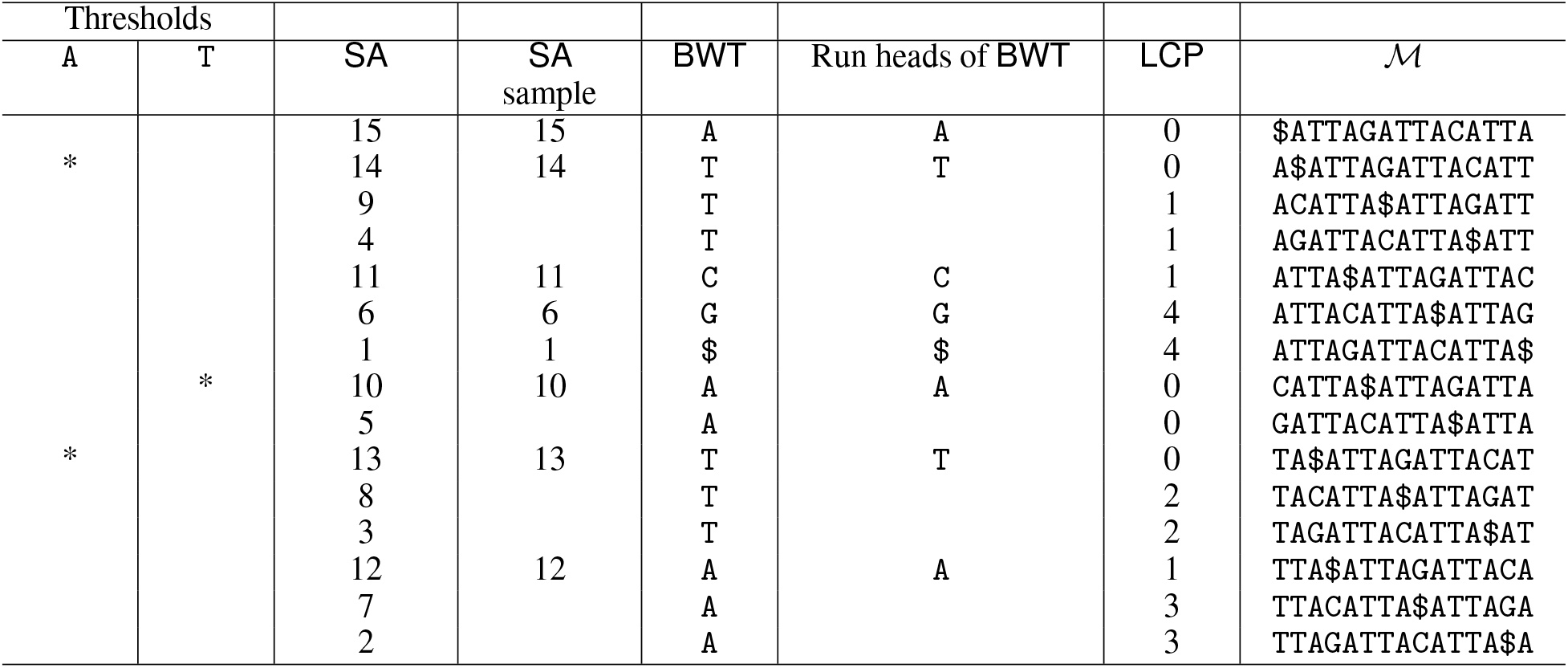
An example of the various components of the *r*-index for the string *S* =ATTAGATTACATTA$, i.e., the SA sample, and RLBWT. The thresholds needed to calculate the matching statistics are also shown. Only the thresholds for A and T are shown because there is only a single run of C, G, and $.

**Figure 1:**
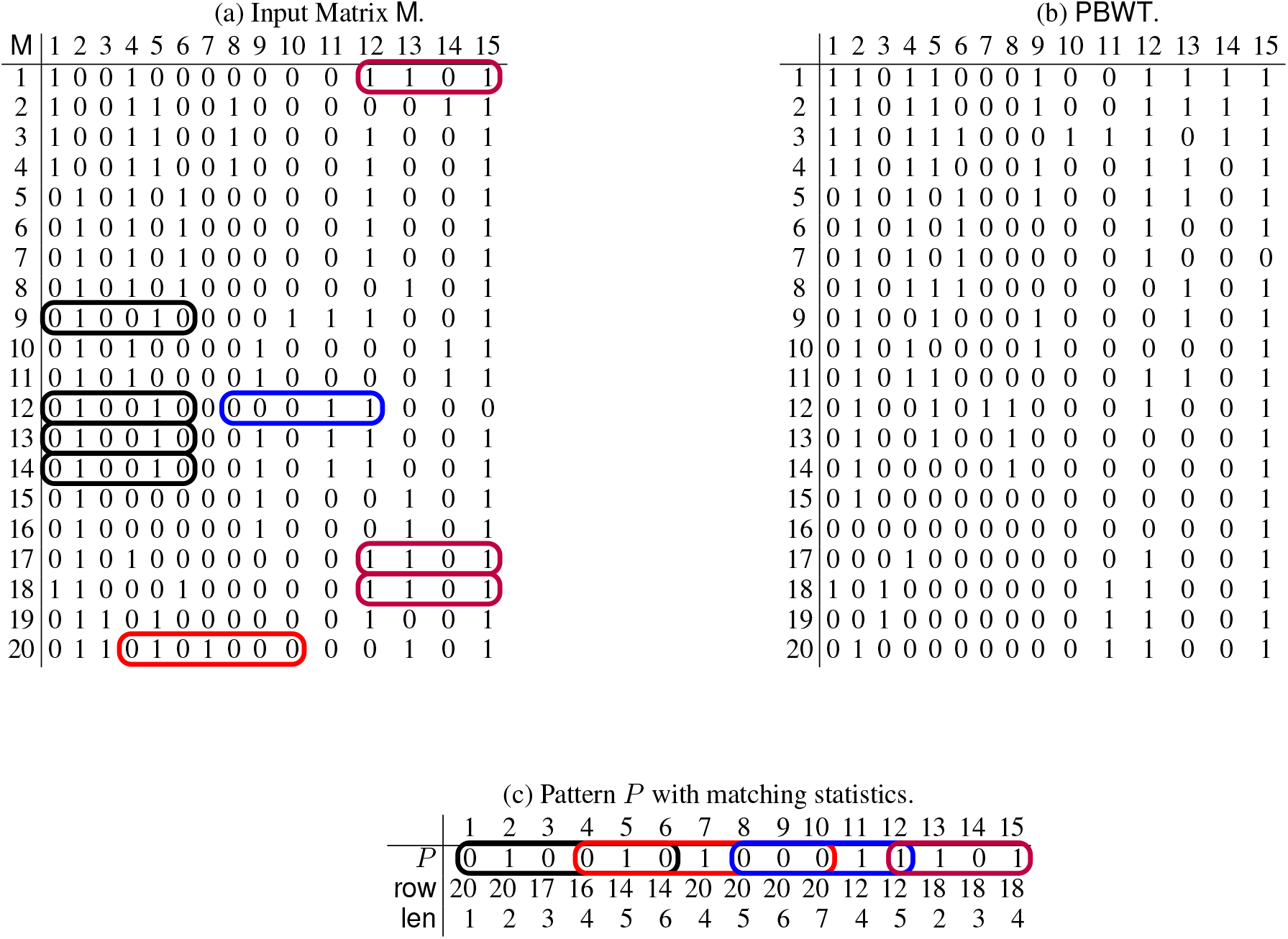
The input matrix M of 20 individuals of 15 bi-allelic sites (a), its PBWT (b), a query pattern *P* and its matching statistics with respect to M (c). SMEMs are circled in both the pattern and the input matrix M.

### 2.9 Straight Line Programs

Aside from BWT, there exist other compression techniques, including Lempel-Ziv and grammar compression. The concept of straight-line programs (SLPs) will be used in our work. SLPs are a representation of the input as a context-free grammar whose language is precisely the input string.

## 3 Definition of SMEMs in the PBWT

In this section, we define SMEMs in the context of the PBWT and describe an approach for finding them. This will allow us to breakdown the algorithm into various steps that can be supported by different data structures.

### 3.1 Definition of the PBWT

Given *h* sequences *S* = {*S*_1_, …, *S*_*h*_}each containing *w* bi-allelic sites corresponding to the same loci, the *j-th prefix array* (PA_*j*_) is the co-lexicographic ordering of the prefixes *S*_1_[1..*j*], …, *S*_*h*_[1..*j*]. The PBWT of M is another *h*×*w* binary matrix, denoted as PBWT[1..*h*][1..*w*], where the first column is the same as the first column of M, and the *j-th* column of PBWT is obtained by stably sorting the rows of M[1..*h*][1..*j* −1] in co-lexicographic order starting at column *j* − 1. More formally, col(PBWT)_1_ = col(M)_1_ and col(PBWT)_*j*_ [*i*] = col(M)_*j*_ [PA_*j* − 1_[*i*]] for all *i* = 1..*h* and *j* = 2..*w*. Figure 1 illustrates an example of the PBWT for an input M. We note that we will frequently use *n* = *h* ·*w* to bound the space- and time-complexity.

The *j-th divergence array* (DA) stores the length of the longest common suffix between the *i*-th and the (*i*−1)-th individuals in the PBWT up to the *j*-th column. More formally, we have DA[1][*j*] = 0 for *j* = 1..*w* and DA[*i*][*j*] = lcs(M[PA_*j*−1_[*i*−1][1..*j*−1]], M[PA_*j*−1_[*i*][1..*j*−1]]) for *i* = 2..*h* and *j* = 1..*w*. We note that in the description of the data structures that follows, it will be convenient to explicitly denote the column for which the divergence array is calculated. For example, if we consider the PBWT shown in Figure 1 and Column 5, then DA[5][7] = 3 because the co-lexicographically 6-th and 7-th row-prefixes (corresponding to 18-th and 16-th rows in the input matrix) up to Column 4 are 0100 and 1100 and their longest common suffix 100 has length 3.

As with the BWT, the PBWT is highly run-length compressible [9]. We denote the PBWT where each column is run-length compressed as RLPBWT. For example, if we store the 15-th column of the PBWT in Figure 1 as (1,6), (0,1), (1,13), where the first number is the bit and the second number is the number of occurrences.

Remembering that we devoted this section to define how to support finding SMEMs in PBWT, we begin by giving a formal definition of a SMEM in the context of the PBWT.

#### Definition 1

*Given w-length input sequences S* = *{S*_1_, …, *S*_*h*_*} (sorted as* M*) and a pattern P* [1..*m*], *we define P* [*i*..*j*], *where* 1 *≤ i ≤ j ≤ w, to be a SMEM if it occurs in one of the input sequences of S and one of the following holds: i) i* = 1 *and j* = *n; ii) i* = 1 *and P* [1..*j* + 1] *does not occur in S; iii) j* = *n and P* [*i −* 1..*n*] *does not occur in S; iv) P* [*i −* 1..*j*] *and P* [*i*..*j* + 1] *do not occur in S*.

## 4 Finding SMEMs in the PBWT

Here, we describe a general algorithm for efficiently finding SMEMs in the input sequences *S* = {*S*_1_, …, *S*_*h*_}using the PBWT.

### 4.1 Matching Statistics in the PBWT

As previously hinted, the problem of finding SMEMs can be cast into the problem of computing matching statistics for *P* using the PBWT, where we slightly adapt the previous definition for our convenience. Given a pattern *P* [1..*m*], the matching statistics of *P* with respect to *S* are an array *A*[1..*w*] of (row, len) pairs such that for each position 1≤*j*≤ *w*: (1) *S*_*A*[*j*]._row[*j*−*A*[*j*].len + 1..*j*] = *P* [*j*−*A*[*j*].len + 1..*j*], and (2) *P* [*j*−*A*[*i*].len..*j*] is not a suffix of *S*_1_[1..*j*], …, *S*_*h*_[1..*j*]. Informally, for each position *j* of the pattern *P, A*[*j*].row is one row of the input matrix M where a longest shared common suffix (of length *A*[*j*].len) ending in position *j* in the pattern *P* and in *S*_*A*[*i*]._row occurs. Observe that the above definition reflects the co-lexicographic ordering in the construction of the PBWT.

We show an example of matching statistics for the input matrix M in Figure 1. SMEMs can be computed from the matching statistics for the PBWT in an analogous manner as to how SMEMs are computed from the matching statistics in the BWT, i.e., we scan the matching statistics from right to left, and report a SMEM at the column *j − A*[*j*].len+ 1 (of the input matrix) of length *A*[*j*].len≥if either *j* = *w*, or *A*[*j*].len ≥ *A*[*j* + 1].len. Informally, *A*[*j*].len≥*A*[*j* + 1].len occurs when we cannot extend to the right the current longest common suffix (of length *A*[*j*].len) shared by *P* and any row in the input matrix. We note that it is fairly straightforward to also report the number of occurrences of a given SMEM in *S*.

### 4.2 Queries Needed to compute matching statistics in the PBWT

Previously, we described how finding SMEMs between a pattern *P* and the given haplotypes is equivalent to computing matching statistics for *P* using the PBWT. Here, we describe how to compute the matching statistics, and in particular, describe that their computation comprises of the application of two specific queries. We introduce these queries and then in the next section describe data structures that can support them. Suppose we have computed *A*[1].row, …, *A*[*j*].row: observe that we need the row index *k* of the PBWT containing *S*_*A*[*j*]._row[*j*]—that is, PBWT[*k*][*j*] = M[PA_*j−*1_[*k*]][*j*] stores *S*_*A*[*j*]._row[*j*]. Now we want to compute *A*[*j* + 1].row and the row index of the PBWT storing *S*_*A*[*j*+1]._row[*j* + 1].

If *P* [*j* +1] = *S*_*A*[*j*]._row[*j* +1], then we can simply set *A*[*j* +1].row = *A*[*j*].row and then compute the row of the PBWT containing *S*_*A*[*j*]._row[*j* + 1] from the number of 0s and 1s in col(PBWT)_*j*_ and the number col(PBWT)_*j*_ .rank_*P* [*j*+1]_(*k*) of copies of *P* [*j* + 1] in col(PBWT)_*j*_ [1..*k*].

If *P* [*j* + 1] ≠ *S*_*A*[*j*]._row[*j* + 1] then we want to choose a row *k*′ of the PBWT such that

- col(PBWT)_*j*_ [*k*′] = *P* [*j* + 1],
- the longest common suffix of 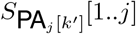 and 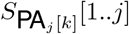 is as long as possible.

If *k*′ < *k* then the length of the longest common suffix of 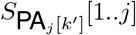 and 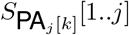 is min(col(DA)_*j*_ [*k*′ + 1..*k*]), and if *k*′ > *k* then it is min(col(DA)_*j*_ [*k* + 1..*k*′]). It follows that we can focus our attention on the last occurrence of *P* [*j* + 1] before PBWT[*k*][*j*] in col(PBWT)_*j*_, and the first occurrence of *P* [*j* + 1] after PBWT[*k*][*j*]. We note these are the last and first copies of *P* [*j* + 1] in two consecutive runs of *P* [*j* + 1] in col(PBWT)_*j*_.

Suppose we have stored the position in *S* of every character at a run boundary in a column of the PBWT. If we also have an LCE data structure for *S*, then we can easily check whether we should set *k*′ to be the row containing the last occurrence of *P* [*j* + 1] before PBWT[*k*][*j*] in col(PBWT)_*j*_, or the row containing the first occurrence after it — and we simultaneously learn *A*[*j* + 1].len.

If we do not have such an LCE data structure but we have a threshold stored that tells us how to choose *k*′—based on whether *k* is above or below the threshold—then we can compute *A*[*j* + 1].row at this point but we do not learn *A*[*j* + 1].len immediately. If we have random access to *S* (or even just to the PBWT), however, then we can still compute *A*[*j* + 1].len with that (although it is more efficient to compute all the row values with one pass and then all the len values with a second one). As we describe later, we can also decide how to choose *k*′ by performing RMQ, PSV and NSV queries on col(DA)_*j*_, and compute *A*[*j* + 1].len if we have random access to col(DA)_*j*_.

If we have also computed the maximal interval [*s, e*] such that 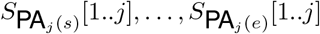 all end with *P* [*j − A*[*j*].len + 1..*j*] and we can perform RMQ, PSV and NSV queries on col(DA)_*j*_, then we can also compute the maximal interval [*s*′, *e*′] such that 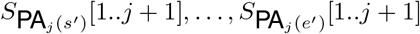 all end with *P* [(*j* + 1) − A[*j* + 1].len + 1..*j* + 1]. Notice *e*′−*s*′ + 1 is thus the number of strings in *S* whose prefixes of length *j* + 1 ends with *P* [1..*j* + 1], which we cannot compute quickly from the matching statistics by themselves. We can break down the computation of matching statistics of a pattern *P* [1..*m*] with respect to *S* via the application of two main queries, which we refer to as the extend query, and the `start query`. To explain further, we let *i, j*∈ [1..*w*] be two integers such that *P* [*i*..*j*] is a suffix of one of *S*_1_[1..*j*], …, *S*_*h*_[1..*j*]. The extend query finds the match of *P* [*i*..*j*] to *P* [*i*..*j* + 1] if and only if *j < w* and *P* [*i*..*j* + 1] is a suffix of one of *S*_1_[1..*j* + 1], …, *S*_*h*_[1..*j* + 1]. The start query finds the smallest integer *i*′∈ [*i*..*j*] such that *P* [*i*′..*j*] is a suffix of one of *S*_1_[1..*j*], …, *S*_*h*_[1..*j*]. Hence, we compute the matching statistics as follows. We assume that we computed the matching statistics up to position *i*′∈ [1..*w*], and use the start query to find the smallest *i′*∈ [*i*..*w*] such that *P* [*i*′..*i*′ + *A*[*i*].len] is a suffix of one of *S*_1_[1..*i*′ + *A*[*i*].len], …, *S*_*h*_[1..*i*′ + *A*[*i*].len]. By minimality of *i*′, we can set *A*[*j*].len = *A*[*j* − 1].len−1 for all *j*∈ [*i* + 1..*i*′ − 1]. Then we find the longest prefix *P* [*i*′..*k*] that is also a suffix of one of *S*_1_[1..*k*], …, *S*_*h*_[1..*k*] using the extend query. We set *A*[*i*′].len = *k*−*i*′ + 1. Since *i*′ > *i*, we can proceed by induction to compute the whole array of matching statistics.

- Referring to the example in Figure 1, we illustrate a section of the execution of the computation of matching statistics and finally of the SMEMs.
- We start from an arbitrary row. In this case, we choose the 20th, where col(PBWT)_1_[20] = 0.
- Since we also have that *P* [1] = 0, we proceed to the next column, and store A[1].row = 20 and A[1].len = 1. To advance by column, we compute the mapping function of row 20 from first column to the second. Observe that the mapping function is used to compute index *k* of col(PBWT)*i*+1 that contains *A*[*i*].row (details in the next section). Hence, the result is that we are mapping on col(PBWT)_2_[15].
- At the second column, we have *P* [2] = col(PBWT)_2_[15], so we can proceed to the next column, storing A[2].row = 20 and A[2].len = 2. Following the mapping of the row 20, we move onto col(PBWT)_3_[19].
- Components
- We have a mismatch at column 3 since *P* [3] ≠ col(PBWT)_3_[19]. At this point, we can move to either the last character of the previous run, col(PBWT)_3_[17], or the first character of the next run, col(PBWT)_3_[20], having PA[17] = 17 and PA[20] = 18. If we look at the input matrix M, we have that, up to column 3 excluded, row 17 has a common suffix to row 20 longer than row 18. So, the best option to maximize the length of the current match is to move to row 17, storing A[3].row = 17 and A[3].len = 3. Now we use row 17 to compute the mapping function from column 3 to column 4.
- We proceed in this way until we complete the computation of A.
- Finally, with a sweep from left to right over A, we can compute all the SMEMs looking at the indices where A[*i*].len *≥* A[*i* + 1].len, as shown in Figure 1 through the colored rounded boxes covering *P*.

## 5 Components: Building Blocks for a PBWT Data Structure

In this section, we present several data structures that support the computation of SMEMs using the PBWT. To accomplish this, we define smaller data structures, which we call components, that can be combined to build data structures that support finding SMEMs in the PBWT. As shown in Figure 2, the components are categorized based on which queries they support, i.e., the `start` or `extend` queries. There are eight components, and six data structures. All six data structures have a mapping structure that allows movement between the columns of the PBWT. The second layer efficiently supports `start` queries, and lastly, the final layer efficiently supports `extend` queries.

**Figure 2:**
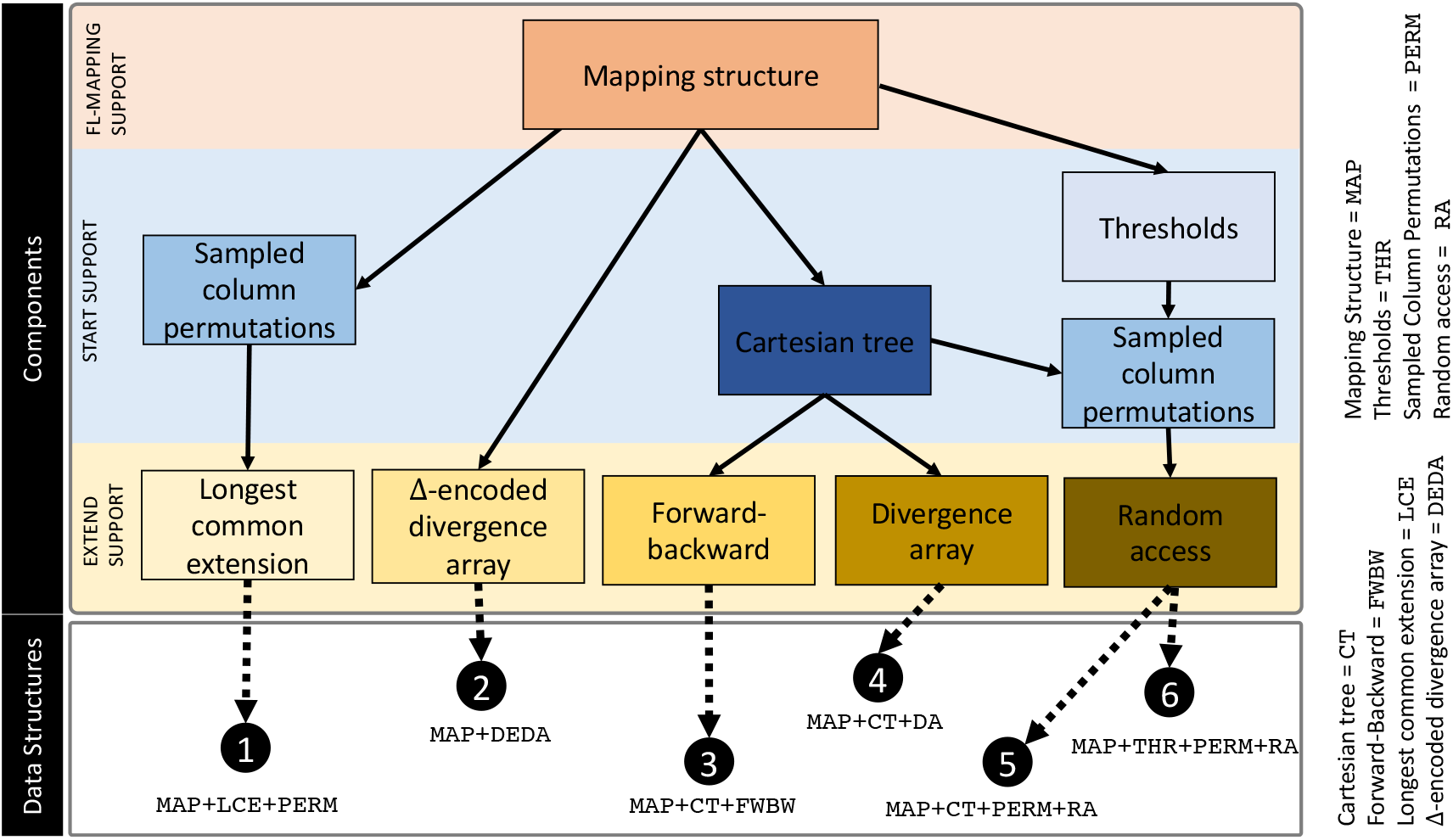
A visualization of our eight data structures supporting finding SMEMs in the PBWT. These data structures are presented as nodes in a tree, and are components of our six solutions for finding SMEMs. Such a solutions is represented by a root-to-leaf path. The various components are described in Section 5 and the data structures are given in Section 6.

### 5.1 Top Level: Mapping Structure

We recall that knowing the location of a specific location of a character, say *S*[*i*], in BWT, we can move backward (i.e., from *S*[*i*] to *S*[*i −* 1]) in the BWT using the LF-mapping as follows:

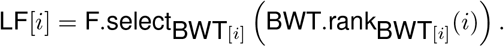

Analogously, all the data structures that we present require a data structure that supports the following query: given the position of a bit in the PBWT, say the *i*-th row and *j*-th column, a mapping structure returns the positions in the PBWT of the bits immediately to the right in M. This is equivalent to forward stepping in the PBWT. Hence, given PBWT[*i*][*j*], we have that the equivalent FL-mapping in the PBWT is as follows. Here, we let *b* = *¬*PBWT[*i*][*j*].

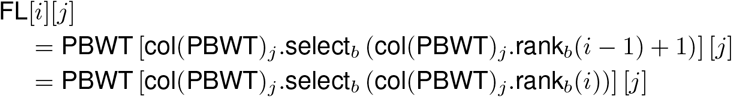

This mapping allows us to step from one column to the next one (to right) in the PBWT. Here, we remind the reader that due to the co-lexicographical ordering on the PBWT, it follows that FL-mapping and forward stepping is the analogous counterparts of the LF-mapping and the backward stepping in the BWT. Similar to the BWT, there exists several mapping structures that can efficiently support FL-mapping in the PBWT.

Three possible data structures that efficiently support mapping are as follows: (1) an uncompressed bitvector occupying roughly *n* bits and answering queries in constant time; (2) a run-length compressed bitvector occupying roughly 𝒪(*r* log(*n/r*)) bits and answering queries in 𝒪(log log *n*)-time; and (3) a simple version of Nishimoto and Tabei’s move structure. See Appendix A for an illustration of (3), which occupies roughly 𝒪(*r* log(*n/r*)) bits and answers queries in constant time [21]. We note that Nishimoto and Tabei’s data structure has yet to be implemented so we implemented the mapping structure as a run-length compressed bitvector.

### 5.2 Second Level: Start Support

In this subsection, we provide a comprehensive discussion of all the data structures that can be used to support `find` queries, i.e., which identify the location of the start of a SMEM.

#### 5.2.1 Sampled Column Permutations

If we use a Cartesian tree but neither the divergence array itself (encoded or unencoded) nor two instances of each component data structure, then it seems we need a way to find at least one occurrence of each SMEM in order to determine its length. We can use the sampled column permutations together with an analogue of Policriti and Prezza’s [24] toehold lemma: for the bits at either end of each run in a column in the PBWT, we store which rows in the input matrix they came from, using a total of roughly 2*r* lg *h* bits; whenever we reach the right end of a SMEM and expand our search interval, the expanded interval must contain the first or last bit in some run of the bits we seek, and we learn from which row of the input matrix it came.

We can use the sampled column permutations and the toehold lemma also with thresholds: Whenever we jump to the first or last bit in a run, we again learn from which row of the input matrix it came. The main difference is that we may jump more than once for each SMEM.

### 5.3 Thresholds

As for the BWT, given the RLPBWT we store for each run in each column (a) the PA sampled at the beginning and at the end of the run and (b) its threshold. Thresholds are the positions of the minimum DA in each run. Formally, let col(PBWT)_*k*_ [*i*..*j*] be a maximal run in the *k*-th column of the PBWT, we store the PA sampled at run boundaries, i.e., the values of PA_*k*_ [*i*], PA_*k*_ [*j*]; and the position of the minimum DA value in the run which is the equivalent of the thresholds for the *r*-index, i.e., RMQDA_[*k*]_(*i, j* + 1), assuming DA[*h* + 1] = 0.

We consider three implementations of the thresholds analogously to those used in Rossi et al. [25]: (1) an uncompressed bitvector occupying roughly *n* bits and answering queries in constant time; (2) a run-length compressed bitvector occupying roughly *r* log(*n/r*) bits and answering queries in 𝒪(log log *n*)-time; and (3) adding roughly *r* lg(*n/r*) bits to the move structure.

#### 5.3.1 Cartesian Trees

If we store a representation of the shape of a Cartesian tree built upon the divergence array, then we can support RMQ, PSV and NSV queries on the divergence array, although those queries return only a position and we cannot support random access easily. We consider three representations of the tree shape: (1) an augmented balanced-parentheses (BP) representation occupying roughly 2*n* + *o*(*n*) bits [10] and answering queries in constant time; (2) a simple DAG-compressed representation (with each non-terminal storing the size of its expansion) answering queries in time bounded by its height; and (3) an interval-tree storing selected intervals corresponding to nodes in the Cartesian tree and answering queries in constant time. We do not include Gawrychowski et al.’s compressed RMQ data structure [14] because we are not aware of an implementation and we see no easy way to estimate its space usage for the divergence array.

The constructions of the BP representation and DAG-compressed representations are standard, but our interval-tree structure needs some explanation. We query the Cartesian tree only while forward stepping through the PBWT and when our search interval contains only 0s and we want a 1, or when it contains only 1s and we want a 0. To proceed, we must ascend the tree and widen our interval (like ascending a prefix tree, discarding the early bits of our pattern) until it contains a copy of the bit we want. Notice our query interval corresponds to a node *v* in the Cartesian tree, and the PBWT interval we seek corresponds to the lowest ancestor *u* of *v* whose interval is not unary. It follows that we need to store in our interval-tree only the PBWT intervals for nodes *u* in the Cartesian tree such that the interval for at least one of *u*’s children is unary but *u*’s interval is not unary.

Since our intervals can nest but not otherwise overlap, we can store our interval-tree in a more space-efficient manner than usual: we write out a string with open-parens, close-parens and 0s, with each open-close pair indicating an interval and the number of 0s before, between and after them indicating its starting point, length and ending point; we encode that string as one bitvector with 0s indicating 0s and 1s indicating parens, and another bitvector with 0s indicating open-parens and 1s indicating close-parens (so the combination of the bitvectors is a wavelet tree for the string); and we store a BP representation of the tree structure of the stored intervals. If we store *k* intervals, then our first bitvector has *n* + 2*k* bits and 2*k* copies of 1, our second bitvector has 2*k* bits with *k* copies of 0 and *k* copies of 1, and the tree structure has *k* nodes and so its BP representation takes 2*k* + *o*(*k*) bits. This means we use roughly 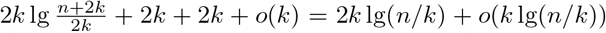 bits, and can answer queries in constant time. We note that, since even our query intervals can nest in but not contain or otherwise overlap any of our stored intervals, we can query with a single endpoint instead of a whole interval.

### 5.4 Third Level: Extend Support

In this subsection, we discuss all data structures that can be combined with the previously described components to support extend queries, that find the lengths of the SMEMs whose start positions were identified using the previous components.

#### 5.4.1 Divergence Array

The simplest possible data structure to support finding the length of each SMEM is to store the uncompressed divergence array. This was originally proposed by Durbin when he described the PBWT. The downside of using this component is the large space requirements—as it would occupy space in bits roughly equal to the sum of the base-2 logarithms of all entries (with 2 added to each entry). As Durbin noted, the divergence array is too big to store easily uncompressed or only loosely compressed: *“Finally, we need a compact representation of [the divergence array]. For now, it is proposed to Huffman encode the differences between adjacent [values in the divergence array] and perhaps only store them for a subset [of columns]. There is probably scope for further improvement here*.*”* In the next subsection, we will discuss storing the divergence array in a compressed format. We note that, although compressing the divergence array using Huffman encoding is possible, it is unknown how this encoding could efficiently support queries.

#### 5.4.2 Δ-encoded Divergence Array

Following Gagie et al. [13], we can build an *O*(*r* log_2_ *n*)-bits balanced SLP for the differentially-encoded (Δ-encoded) divergence array. That is, we store each entry of DA[*i*][*j*] with *i >* 1 as the difference DA[*i*][*j*] *−* DA[*i −* 1][*j*]; for *i* = 0 we always have DA[*i*][*j*] = 0. If PBWT[*i*..*i* + *𝓁 −* 1][*j*] is a run of equal bits in the *j*-th column of the PBWT and col(PBWT)_*j*+1_[*i*′..*i*′ + *𝓁 −* 1] are the bits immediately to their right in the input matrix, then DA[*i*′ + *k*][*j* + 1] = DA[*i* + *k*][*j*] + 1 for 1 *≤ k ≤ 𝓁 −* 1. Therefore,

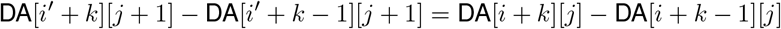

for 1≤*k*≤*𝓁*−1, so the Δ-encoding of DA[*i*′ + 1..*i*′ + *𝓁*−1][*j* + 1] is the same as that of DA[*i* + 1..*i* + *𝓁*−1][*j*]. It follows that if the PBWT is highly run-length compressible, then the Δ-encoded divergence array is highly repetitive.

This property is similar to the Δ-encoded LCP array, which has a string attractor [16] of size 𝒪 (*r*) [13, Lemma 6.2] and thus, it can be represented as a straight-line program (SLP) occupying 𝒪 (*r* log_2_ *n*) bits [16, Lemma 3.14]. Thus, we can follow the same steps of Gagie et al. [13, Lemma 6.2] to conclude that we can also find an SLP for the linearized Δ-encoded divergence array in the same amount of space.

Increasing the size of this SLP by a small constant factor, we can store at each non-terminal the length, sum and minimum prefix sum of its expansion, and thus support random access, RMQ, PSV, and NSV queries on the divergence array in 𝒪 (log *n*)-time. To see how this works, suppose we want to access DA[*i*][*j*], which is the *t*th entry in the linearized divergence array, where *t* = *ih* + *j*. By the definition of the Δ-encoding, DA[*i*][*j*] is equal to the sum of the first *t* values in the linearized Δ-encoded divergence array, so we can compute it much like we would compute a partial sum of keys in a binary search tree.

To compute the partial sum equal to *DA*[*i*][*j*], we start at the root of the SLP’s parse tree and descend to its *t*th leaf (which we can find in time proportional to its depth, because we have the sizes of the non-terminals’ expansions stored). As we descend the path, we sum all the first *t*−1 numbers in the Δ-encoded divergence array: these numbers are in the expansions of symbols not on the path, that are left children of symbols on the path, and we have those left children’s sums stored. Finally, we add the *t*th number in the Δ-encoded divergence array, which is stored at the *t*th leaf.

For RMQs, we use the fact that the sum of the Δ-encoded entries to the left of a non-terminal’s expansion, plus the minimum prefix sum of its expansion (of Δ-encoded entries), is the minimum entry in the corresponding interval of the unencoded divergence array. Given a query range [*i*..*j*], we can find 𝒪 (log *n*) subtrees of the parse tree whose leaves are the *i*-th through *j*-th, and return the minimum unencoded divergence entry in that interval. For PSV queries, we descend to the specified *t*-th leaf of the parse tree and re-ascend until we find a left child *v* hanging of the path we descended, whose expansion includes a smaller divergence-array value than the *t*-th. We then descend into and find the rightmost entry in *v*’s expansion smaller than the *t*-th entry. NSV queries are symmetric.

#### 5.4.3 Longest Common Extension

Next, we consider the addition of an LCE data structure. Suppose we have arrived at column *j* + 1 and we know that the longest suffix of pattern *P* [1..*j*] that occurs in M ending at column *j* has an occurrence immediately followed by PBWT[*i*][*j* + 1], and we know which row of M that bit PBWT[*i*][*j* + 1] comes from, but *P* [*j* + 1]*≠* PBWT[*i*][*j*].

By the definition of the PBWT, there is an occurrence of the longest suffix of *P* [1..*j*] that occurs in M ending at column *j*, ending at either:

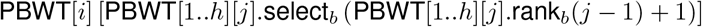

or

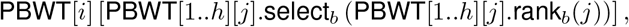

where *b* = *¬*PBWT[*i*][*j*] = *P* [*j*].

We can efficiently choose the former or the latter if we have a data structure supporting LCE queries on the input matrix (backwards) and we have stored the rows in M from which these two bits come. For more details, we refer the reader to [4].

Although there are many compressed LCE data structures, a large fraction has not been implemented. Here, we list two practical implementations:

- an SLP of the input matrix (read row-wise) answering queries in *O*(log *n*) time,
- a plain representation of the matrix (with 8 bits packed into each byte) occupying roughly *n* bits and answering queries in time proportional to the length of the LCE. In particular, allowing word-packing with a word size of Θ(log *n*), we can answer an LCE of length *𝓁* in *O*(1 + *𝓁/* log *n*)-time.

#### 5.4.4 Forward-Backward

Suppose we use a Cartesian tree to maintain the invariant that our search interval in the PBWT contains all the bits immediately following occurrences of the longest suffix of the prefix of the pattern that we have processed so far, that occur in the desired columns of the PBWT. If that search interval contains a copy of the next bit of the pattern, then we proceed by forward stepping, without consulting the Carteisan trees.

The only time we query a Cartesian tree is when the search interval does not contain a copy of the next bit of the pattern—meaning we have reached the right end of a SMEM. It follows that, using the mapping structure and the Cartesian trees, we can find the right endpoints of all the SMEMs. If we keep instances of all our components for the reversed input matrix, we can also find all the left endpoints of the SMEMs. Since SMEMs are maximal, they cannot nest, so we can easily pair up the endpoints and obtain the SMEMs. This roughly doubles the time and space.

#### 5.4.5 Random Access

Lastly, the simplest possible component is a compressed data structure of the original input that provides efficient random access to the input. Trivially, we can see that such a data structure can be used to find the length of a given SMEM. We note that although the total length of the SMEMs can be quadratic in the length of the pattern, the fact they cannot nest implies we need only a linear number of random accesses. In fact, if we combine a random access data structure with Cartesian trees then the number of random accesses is equal to the number of SMEMs, and the total length of the sequence that we extract from M is linear in the length of the SMEMs. There are many data structures that support random access to the input matrix M, two notable ones are (a) an SLP of M (read row-wise) answering queries in 𝒪 (log *n*) time, and (b) a plain representation of M (with 8 bits packed into each byte) occupying roughly *n* bits and allowing access to each bit in constant time.

## 6 Composite Data Structures for the PBWT

In this section, we combine the various components that we described in the previous section to create data structures that efficiently support finding SMEMs in the PBWT. Due to brevity and ease of description, we already described some of these in the previous section when we described the components. More specifically, the “Mapping Structure + Cartesian Tree + Forward-backward” and “Mapping Structure + Cartesian Tree + Sample Column Permutations + Random Access” were described in the previous section. In what follows, we describe the remaining four compositions. Thus, we will have a total of six data structures for the PBWT.

### 6.1 Using the Divergence Array

The basic idea is already sketched in the case for Cartesian trees in Section 5.3.1. The idea is, given a search interval *I* that contains only 0 bits, and we want a 1 bit (the other way follows by symmetry), we query for the minimum value *v* in the divergence array, and compute the positions *a* and *b* of the previous and next smaller value of *v*, respectively. These positions obey the invariant that *I*⊂(*a*..*b*) and (*a*..*b*) matches a node in the Cartesian tree. Finally, having access support to the divergence array allows us to determine the length of a match. To obtain RMQ, NSV, and PSV support, we can use the Δ-encoded divergence array (Section 5.4.2) or a Cartesian tree. Hence, we obtain the solutions “Mapping Structure + Δ-Encoded Divergence Array” and “Mapping Structure + Cartesian Tree + Divergence Array”.

### 6.2 Mapping Structure + Longest Common Extensions + Sampled Column Permutations

Similarly to what is described in Boucher et al. [4], we can compute the matching statistics in a single-pass algorithm over the input pattern *P*. We process the pattern *P* from left to right, and at each step, we use the LCE data structure to compute the length of the longest common suffix between the longest match of the pattern at the current position, and the sequences in the matrix, to decide which candidate has the longest match with the pattern. We assume that we computed the matching statistics component up to position *k*− 1, and are processing the *k*-th column. Let *i* be the row of the PBWT that matches the longest suffix of *P* [1..*k −* 1] that is suffix of *S*_1_[1..*k −* 1], …, *S*_*h*_[1..*k −* 1], and let *p* be the corresponding row in M i.e., for all 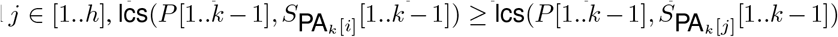 with *p* = PA_*k*_ [*i*]. If col(PBWT)_*k*_ [*i*] = *P* [*k*] then row *i* can be used to extend the match, hence we can assign *A*[*k*].row = *p, A*[*k*].len = *A*[*k*−1].len+1, *i* = LF(*i, k*), and *p* does not change. Otherwise, if col(PBWT)_*k*_ [*i*] ≠ *P* [*k*], let col(PBWT)_*k*_ [*s*..*e*] be a maximal run containing position *i*, then the longest suffix of *P* [1..*k*] that is suffix of *S*_1_[1..*k*], …, *S*_*h*_[1..*k*] is either the one corresponding to the preceding end or following start of a run of *P* [*k*] in col(PBWT)_*k*_ with respect to position *i*, i.e., either 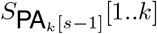 if *s* > 1 or 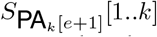 if *e < n*. Since for each run we have stored the samples of PA at the beginning and at the end of each run, and we have the value of *p*, we can use the LCE data structure to compute the two longest common suffixes above, keep the longest, and continue matching from there.

### 6.3 Mapping Structure + Thresholds + Sampled Column Permutations + Random Access

Similarly to what is described in Rossi et al. [25], we can compute the matching statistics in a two-pass algorithm over the input pattern *P*. During the first scan, we process the pattern *P* from left to right, storing for each position the row component of the matching statistics. In the second scan, we process the pattern *P* from right to left, and with the use of a random access data structure on the binary array M, we compute the len component of the matching statistics.

We proceed as in the previous case, however, instead of using the LCE data structure to decide which candidate to choose, we use the thresholds. Let *t* be the position of the threshold in the current run. By the definition of the thresholds, if the position *i* < t then 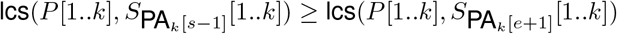 hence we can assign *A*[*k*].row = *p* = PA_*k*_ [*s−* 1] and *i* = LF(*s−* 1, *k*). Otherwise, 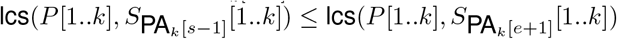 hence we can assign *A*[*k*].row = *p* = PA_*k*_ [*e* + 1] and *i* = LF(*e* + 1, *k*).

Once we have collected all the occurrences of maximal matches between the pattern and the matrix, we can compute the lengths of those matches in linear time by scanning the pattern *P* from right to left. Let us assume that we have computed the lengths up to position *i* + 1, and we want to compute the length of position *i*. Since it holds that *A*[*i*].len ≥ A[*i* + 1].len − 1, we can compute the length *A*[*i*].len by comparing the characters in the pattern *P* and in the matrix in correspondence of row *A*[*i*].row and the column *i* − A[*i* + 1].len − 1.

## 7 Experiments and Discussion

In this section, we provide experimental evaluations of our presented data structures. We begin by benchmarking the memory usage of each of our components, leading to an estimation of the memory-efficiency of our six data structures. Based on these experiments, we fully implement two of these data structures and show the scalability of them on real data.

### 7.1 Benchmarking for Data Structure Selection

#### Experimental set-up

For our experiments, we replicate the simulated dataset used in Durbin’s original PBWT experiments [9], covering a 20-megabases sequence from 100,000 haplotypes. That is, we run the sequentially Markovian coalescent simulator MaCS of Chen et al. [6] with command line parameters 100000 2e7 -t 0.001 -r This generates a haplotype matrix with 100,000 individuals (rows) and 360,000 variation sites (columns). The matrix is stored in transposed form to allow efficient streaming of the columns from disk.

Next, we subsampled the dataset with a parameter *ξ* ∈{0.01, 0.03, 0.05, 0.08, 0.10} such that, given a column of length *h* having *o* ones, we skip this column if *o/h < ξ*. Durbin used *ξ* = 0.05 and further subsampled only 10% of that dataset, or about six thousand columns. We consider the other values of *ξ* because, as they rise, the datasets become less and less repetitive. We thus gain an insight into how much different combinations of component data structures depend on repetitions.

All benchmarking experiments were ran with an Intel Core i3-9100 CPU (3.60GHz) with 128 GB RAM, running Debian 11.

#### Implementation

All our methods were implemented in C/C++. The mapping structure was implemented using run-length compressed bit vectors. The differentially encoded div-array was implemented with grammar compression. The Cartesian tree was also implemented with grammar compression for the dataset with density 0.1 and the interval representation for the rest. Forward and Backward was implemented by building the data structures in both the forward and backward directions. The sampled column permutations were obtained by sampling at run boundaries. The longest common extension query data structure and the random access was implemented with a grammar built on the input matrix. Lastly, thresholds were implemented as a sparse bit vector.

Next, we consider the combination of the data structure. Each combination uses a mapping structure as basis. We denote the resulting data structures based on the other components as follows (cf. Figure 2):

1. LCE data structure + sampled column permutations by `MAP+LCE+PERM;` and
2. Δ-encoded divergence array by `MAP+DEDA;`
3. Cartesian tree + forward-backward by `MAP+CT+FWBW;`
4. Cartesian tree + divergence array by `MAP+CT+DA;`
5. Cartesian tree + sampled column permutations + random access by MAP+CT+PERM+RA;
6. thresholds + sampled column permutations + random access by MAP+THR+PERM+RA.

Our code is available at https://github.com/koeppl/pbwt and https://github.com/dlcgold/rlpbwt and the simulated haplotype matrix is available at http://dolomit.cs.tu-dortmund.de/tudocomp/pbwt_matrix.xz.

### Benchmarking the memory usage

Table 2 shows the statistics of the datasets from which we estimate our component data structures’ sizes. Since it is hard to predict the size of SLPs, we actually built them for the Δ-encoded divergence arrays, the DAG-compressed Cartesian trees and the input matrices. For the Cartesian trees and input matrices, we tried both BigRePair [12] and SOLCA [30], and then applied Shaped_SLP encoding [11], but for the Δ-compressed divergence arrays we were able to run only BigRePair. (Besides other RePair implementations, it is currently the only grammar software we are aware of that supports input alphabets larger than the byte alphabet. It is not, however, guaranteed to produce balanced grammars, thus undermining our query-time bounds.)

**Table 2:**
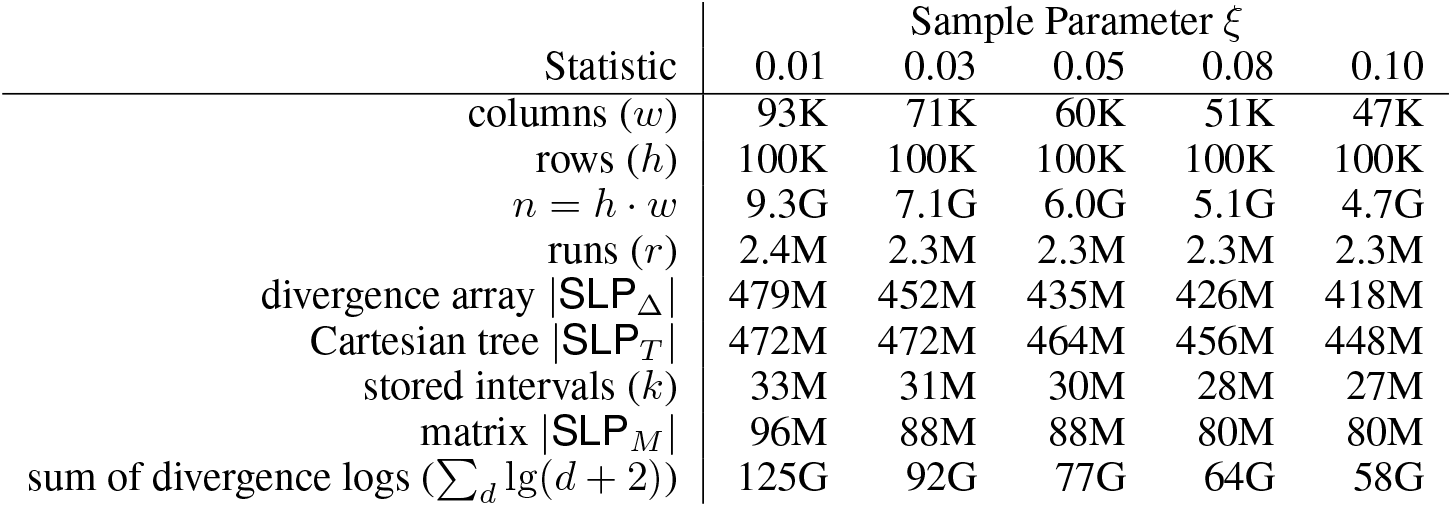
Statistics for the datasets. The size of each SLP is given in bits for consistency with how the other statistics are used to estimate sizes. K, M and G mean thousands, millions and billions, respectively. Notice that many statistics shrink as the sample parameter rises, but only because the number of columns decreases more rapidly.

Table 3 shows the formulas we used to estimate the sizes in bits of our component data structures (repeated from Section 5), and those estimates for each dataset. Although we computed actual SLPs, we did not fully augment them, and thus those sizes should also be taken as estimates. For most of the components, one implementation is the clear winner, but the size of the Cartesian tree implemented with an SLP is smaller for the first two datasets, and the estimate of the implementations with intervals is smaller for the last three datasets.

**Table 3:** The estimated size in bits of each of our component data structures for each of the datasets. In most cases one implementation is the clear winner, but the size SLP_*T*_ of the Cartesian tree implemented with an SLP is smaller for the first two datasets, and the estimate 2*k* lg(*n/k*) of the implementations with intervals is smaller for the last three datasets. M denotes megabytes and G denotes gigabytes. (We do not list Forward-Backward here as it is not a new data structure, only two instances of the Mapping Structure and the Cartesian Tree.)

| Component | Estimate Formula | Sample Parameter $\xi$ | | | | |
| --- | --- | --- | --- | --- | --- | --- |
|  |  | 0.01 | 0.03 | 0.05 | 0.08 | 0.10 |
| Mapping structure | $\min(n, 2r \lg(n/r))$ | 57M | 53M | 52M | 51M | 51M |
| Δ-encoded divergence array | $\vert \text{SLP}_\Delta\vert $ | 479M | 452M | 435M | 426M | 418M |
| Cartesian tree | $\min(2n, \vert \text{SLP}_T\vert , 2k \lg(n/k))$ | 472M | 472M | 458M | 420M | 402M |
| Longest common extension | $\min(\vert \text{SLP}_M\vert , n)$ | 96M | 88M | 88M | 80M | 80M |
| Thresholds | $2r \lg(n/r)$ | 57M | 53M | 52M | 51M | 51M |
| Sampled column permutations | $2r \lg h$ | 80M | 76M | 76M | 76M | 76M |
| Divergence array | $\sum_d \lg(d+2)$ | 125G | 92G | 77G | 64G | 58G |
| Random access | $\min(\vert \text{SLP}_M\vert , n)$ | 96M | 88M | 88M | 80M | 80M |

Table 4 shows the total estimated sizes of the various combinations that can be used to find SMEMs. The combination that yielded the most efficient space usage was `MAP+LCE+PERM`, which is analogous to PHONI [4] in the context of the BWT. MAP+THR+PERM+RA was a very close second, which is analogous to the *r*-index implementation of Rossi et al. Finally, `MAP+DEDA` performed surprisingly well. It is worth noting that that combination was inspired by a similar structure suggested by Gagie et al. [13] but not implemented because it was thought to be too impractical. Of course, augmenting the SLP will increase its size, but a careful implementation should still be competitive.

**Table 4:** The estimated size in bits of the combinations of component data structures (choosing the smallest implementations) that support SMEM-finding in the PBWT. M denotes megabytes and G denotes gigabytes. In boldface the best performance.

| Data Structure | Sample Parameter $\xi$ | | | | |
| --- | --- | --- | --- | --- | --- |
|  | 0.01 | 0.03 | 0.05 | 0.08 | 0.10 |
| (1) MAP + LCE + PERM | **233M** | **217M** | **216M** | **207M** | **207M** |
| (2) MAP + DEDA | 536M | 505M | 487M | 477M | 469M |
| (3) MAP + CT + FWBW | 1.1G | 1.1G | 1.0G | 942M | 906M |
| (4) MAP + CT + DA | 126G | 93G | 78G | 64G | 58G |
| (5) MAP + CT + PERM + RA | 705M | 689M | 674M | 627M | 609M |
| (6) MAP + THR + PERM + RA | 290M | 270M | 268M | 258M | 258M |

The sizes listed in Table 4 are absolute, and so decrease as *ξ* increases even though the datasets become less repetitive, because fewer columns are selected and the datasets shrink significantly. As expected, as *ξ* grows, the compression suffers for all the methods but `MAP+LCE+PERM` and MAP+THR+PERM+RA maintain their lead. For applications requiring counts of how often each SMEM occurs, however, the Δ-encoded divergence array or a Cartesian tree could still be useful.

### 7.2 Results on PBWT Data Structures

#### Experimental set-up

Based on our benchmarking in the previous section, we selected two data structures that were deemed to have superior performance and evaluated them on the 1000 Genomes Project data—namely, `MAP+LCE+PERM` and MAP+THR+PERM+RA. To accomplish this, we downloaded the VCF files for the 1000 genomes project data and then converted the VCF files to bi-allelic. This latter step can be accomplished trivially using various methods. We used `bcftools view -m2 -M2 -v snps` [8]. Then we considered increasingly larger datasets from the converted VCF files and measured the time and space for building the data structures and then the query time, more precisely the performance in computing SMEMs. All these tests were performed without applying a multithreading approach.

From the 1000 Genome Project dataset, We selected for their size the panels related to Chromosomes 22, 20, 18 and 16, having 5008 samples and a number of bi-allelic sites (selected by bcftools) increasing from ∼1 million to ∼2.5 millions. In addition we tested the data of Chromosome 1, being one of the largest panels available (~6 millions of bi-allelic sites). The data used are available at https://ftp.1000genomes.ebi.ac.uk/vol1/ftp/release/20130502/. Experimentally, we observed these panels are sparse, having indeed few ‘1’s compared to ‘0’s. The sparsity of data is also confirmed by the average number 12 of runs per column in the run-length encoded PBWT.

Regarding tests on the 1000 Genomes Project data, we ran them on a machine with an Intel Xeon CPU E5-2640 v4 (2.40GHz) with 756 GB RAM and 768 GB of swap, running Ubuntu 20.04.4 LTS.

#### Results on 1000 Genomes Project data

We decided to compare the two data structures `MAP+LCE+PERM` and MAP+THR+PERM+RA with the original PBWT implementation of Durbin, available at https://github.com/richarddurbin/pbwt. In detail, we ran the `matchIndexed` algorithm, corresponding to **algorithm 5** of the original PBWT paper [9], and the `matchDynamic` algorithm, corresponding to the one named “batch” in the results section of the same paper. In these tests the SLP is already available when needed but it must be specified that its construction takes a long time, up to 4 hours in the case of the Chromosome 1 panel.

Figures 3 (a) and (b) show respectively construction maximum memory usage and time. Note that the PBWT does not build all indices needed for mapping during the construction, while run-length encoded PBWT builds all the structures necessary for the computation of SMEMs. MAP+THR+PERM+RA, with plain representation of the matrix, requires more memory and time construction having to build such representation.

**Figure 3:**
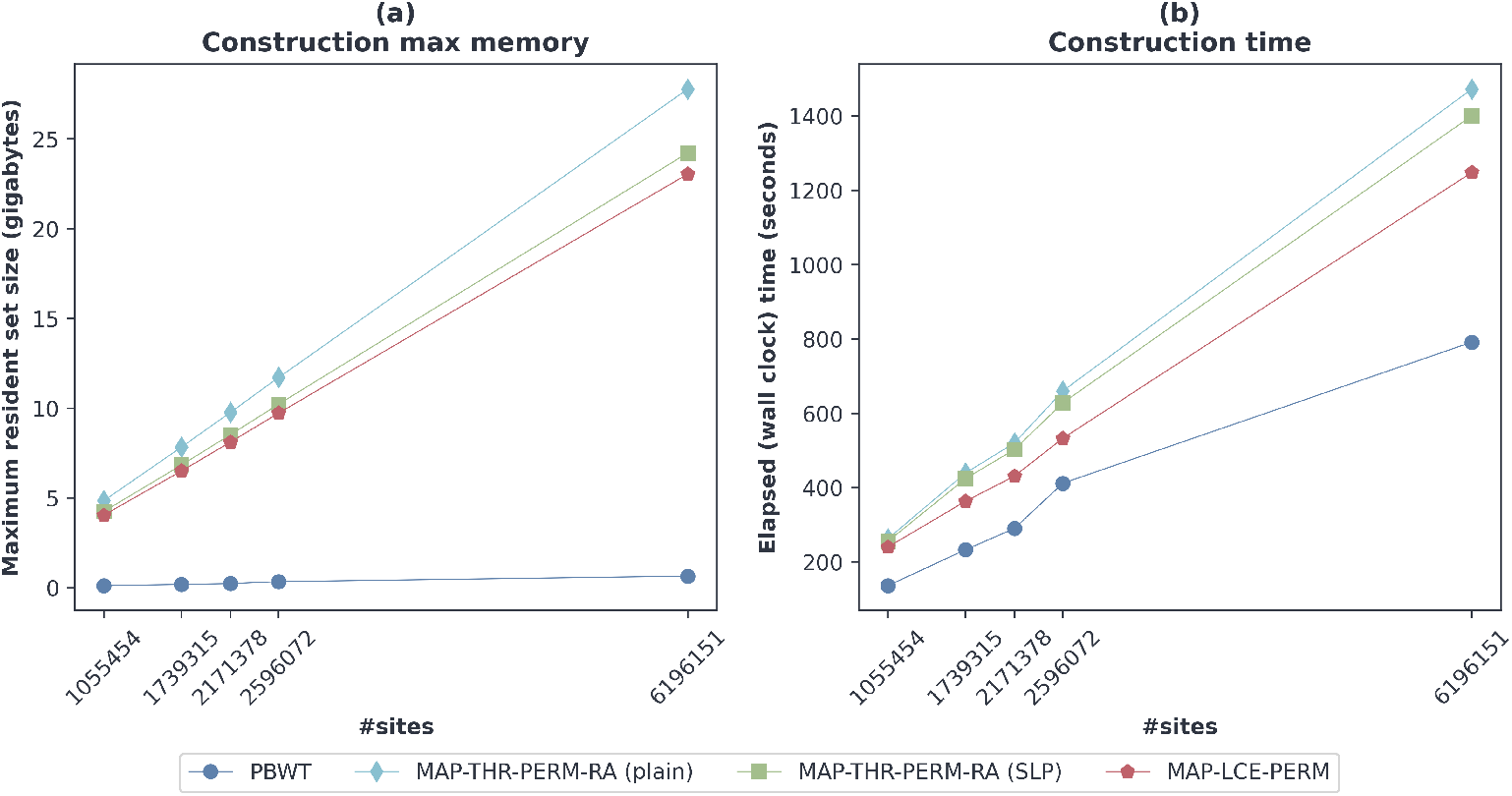
Memory (a) and time (b) to build the run-length encoded PBWT and PBWT. All the panels have 4908 samples.

In order to test the performance of computing SMEMs, 100 sequences were extracted from the input panels, to use them as queries. Thus, the tested panels have 4908 samples. Figure 4 (a) and (b) show respectively the maximum memory usage and the time of the SMEMs computation. These results include data structure loading and any other computation required by Durbin’s PBWT implementation (such as populating the various arrays for `PBWT Indexed`). `PBWT Dynamic` has the best performances when running multiple queries. Regarding maximum memory usage, all the run-length encoded solutions tested perform better than `PBWT Indexed;` the latter requires up to 20 times the memory required by the variants of the run-length encoded PBWT. `MAP+THR+PERM+RA (plain)` has lower execution times thanks to the fact that it makes random access to the panel in constant time, requiring more memory. MAP+THR+PERM+RA (SLP) has longer execution times than `MAP+LCE+PERM` due to the length of the panel and the queries, which require a large number of accesses to the SLP. For each panel, we obtained the following average numbers of SMEMs per 100 queries: 1184 SMEMs for chromosome 22 (1055454 sites), 1416 SMEMs for chromosome 20 (1739315 sites), 1708 SMEMs for chromosome 18 (2171378 sites), 2281 SMEMs chromosome 16 (2596072 sites) and 4953 SMEMs for chromosome 1 (6196151 sites).

**Figure 4:**
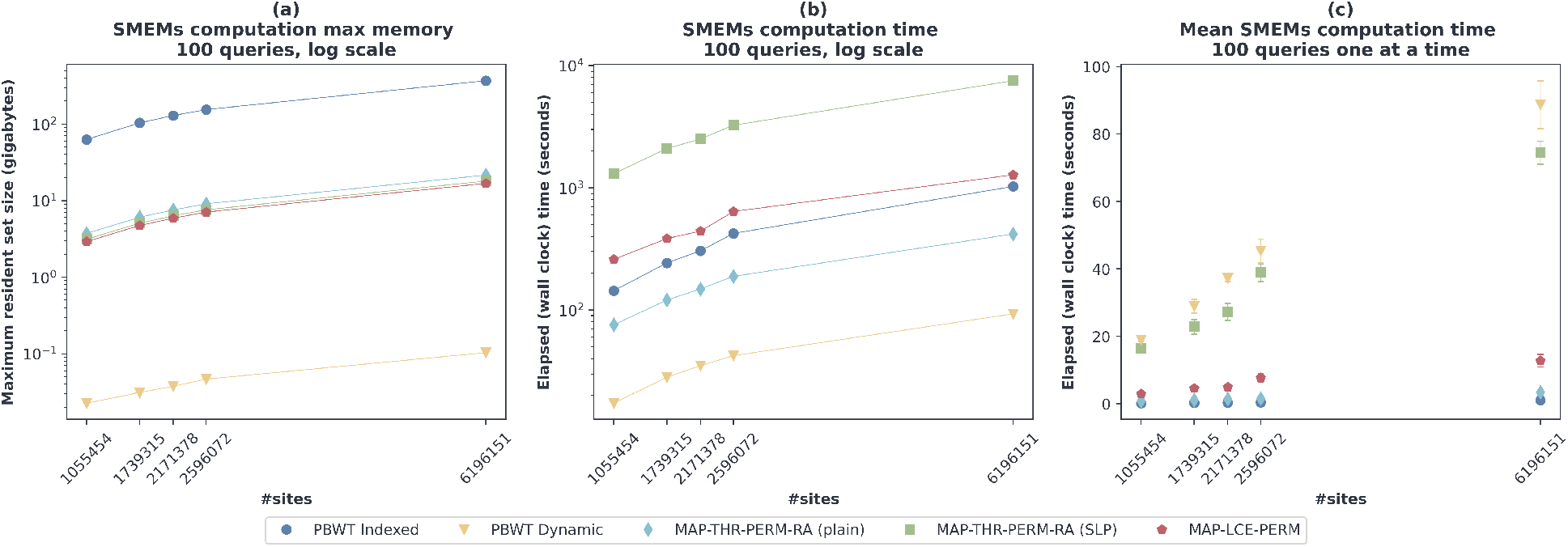
Memory (a), Time (b) and mean time for one query at a time (c) to compute SMEMs with 100 queries. All the panels have 4908 samples and in (c) error bars represent the standard deviation.

Figure 4 (c) shows mean time and standard deviation of the computation of SMEMs with a single query, obtained by executing 100 queries one at a time. In this case, the query functions were tested without considering the loading of structures or any other computation required by Durbin’s PBWT implementation. `PBWT Dynamic` has the worst performance in time due to the fact that it is optimized for handling multiple queries, using an algorithm that studies also the PBWT of the queries panel (whose construction time is not counted in the figure). As expected, concerning run-length encoded PBWT variants, `MAP+THR+PERM+RA (plain)` has the best time performances, comparable to those of `PBWT Indexed`, but with the reduced memory usage discussed earlier, having that `PBWT Indexed`, according to Durbin, requires 13*n* bytes that are independent of the number of queries. The results of the other run-length encoded PBWT variants are consistent with those discussed above.

## 8 Conclusion

In this paper, we consider the problem of computing SMEMs in the PBWT in order to solve Durbin’s problem of finding the maximal haplotype matches. The PBWT has been comparatively overlooked by the data structures community, which motivates us to consider the space-efficiency of various approaches. We not only give a theoretical bound to the space complexity, but also partially implement several of the approaches to estimate their performance in practice. It seems the approaches following Rossi et al. [25] are likely to be the smallest, but using an SLP for a Δ-encoded divergence array may be competitive, especially when counts are desired. This suggests that similar techniques may work on BWTs of pangenomes, which are much larger but also much more repetitive than the datasets we have considered here.

## 9 Acknowledgments

The authors thank Jarno Alanko for helping us with the experiments, Yakov Nekrich for the interval-tree structure for the Cartesian Trees, and the anonymous reviewers for their valuable suggestions.

## A Appendix: Move Structure for the PBWT

Nishimoto and Tabei [21] recently showed how to avoid the use of rank queries while performing the LF mapping in the *r*-index (or backward searches, when the alphabet is polylogarithmic). Their result was surprising but in retrospect it should have been obvious for the PBWT.

Suppose that, for each run PBWT[*i*..*i* + *𝓁*−1][*j*] in the PBWT, we store the following: (a) *i*; (b) *i*′ such that PBWT[*i*′][*j* + 1] is immediately to the right of PBWT[*i*][*j*] in the input matrix; and (c) the index *x* of the run in column *j* + 1 that contains PBWT[*i*′][*j* + 1]. For each column of PBWT, we store a list of these triples (*i, i*′, *x*) sorted by the first component. Figure 5 shows the last 12 columns of the PBWT of Durbin’s example matrix, and Table 6 shows the corresponding tables of triples.

**Figure 5:**
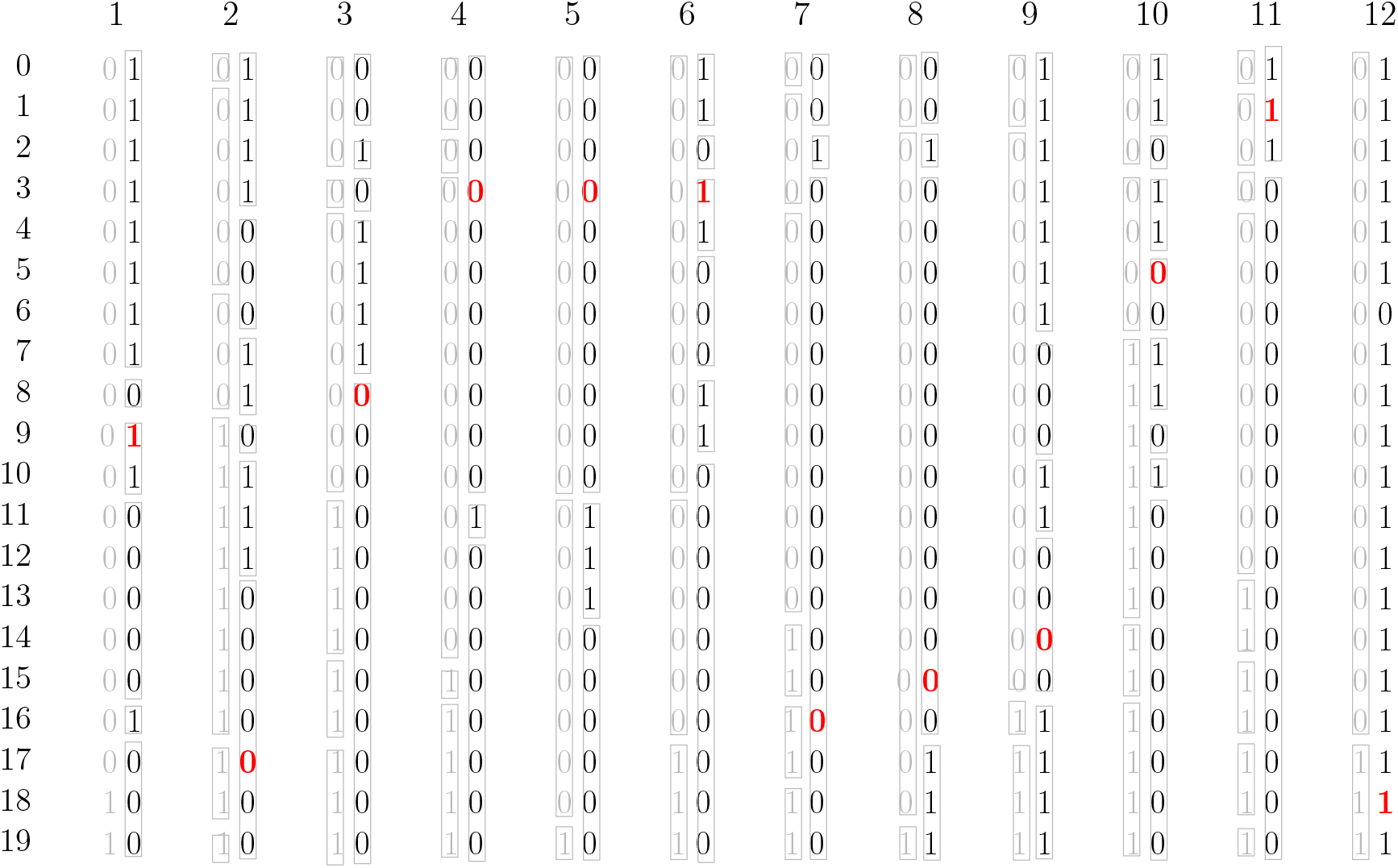
The last 12 columns of the PBWT for Durbin’s example matrix. Each column entry is shown in black except for the bit from a particular row of that original matrix, which is red, with the sorted bits of the previous column to the left in grey. The grey boxes show how the bits in the runs in each column stay together when the column is stably sorted.

Suppose we know initially that the first red bit in Figure 5 is in row 9 and run 3 of the first column, and we want to extract the other red bits (which come later in the same row of the input matrix). If we know the *t*th red bit is in row *i* of the *t*th column, and the triple for its run is (*x, y, z*), then the (*t* + 1)st red bit is in row *i*′ = *y* + *i*−*x*. To find the triple for its run, we start at triple *z* in the (*t* + 1)st table and scroll down until we find the last triple (*x*′, *y*′, *z*′) such that *x*′≤ *i*′. In other words, we can replace rank queries by table lookup, at the cost of sometimes having to scroll down the tables (but usually not very far in practice [5]). In Figure 6, the triples in red contain the information we need to extract the red bits in Figure 5; where they differ from the triples at which we first arrive by table lookup, the latter are shown in blue.

**Figure 6:**
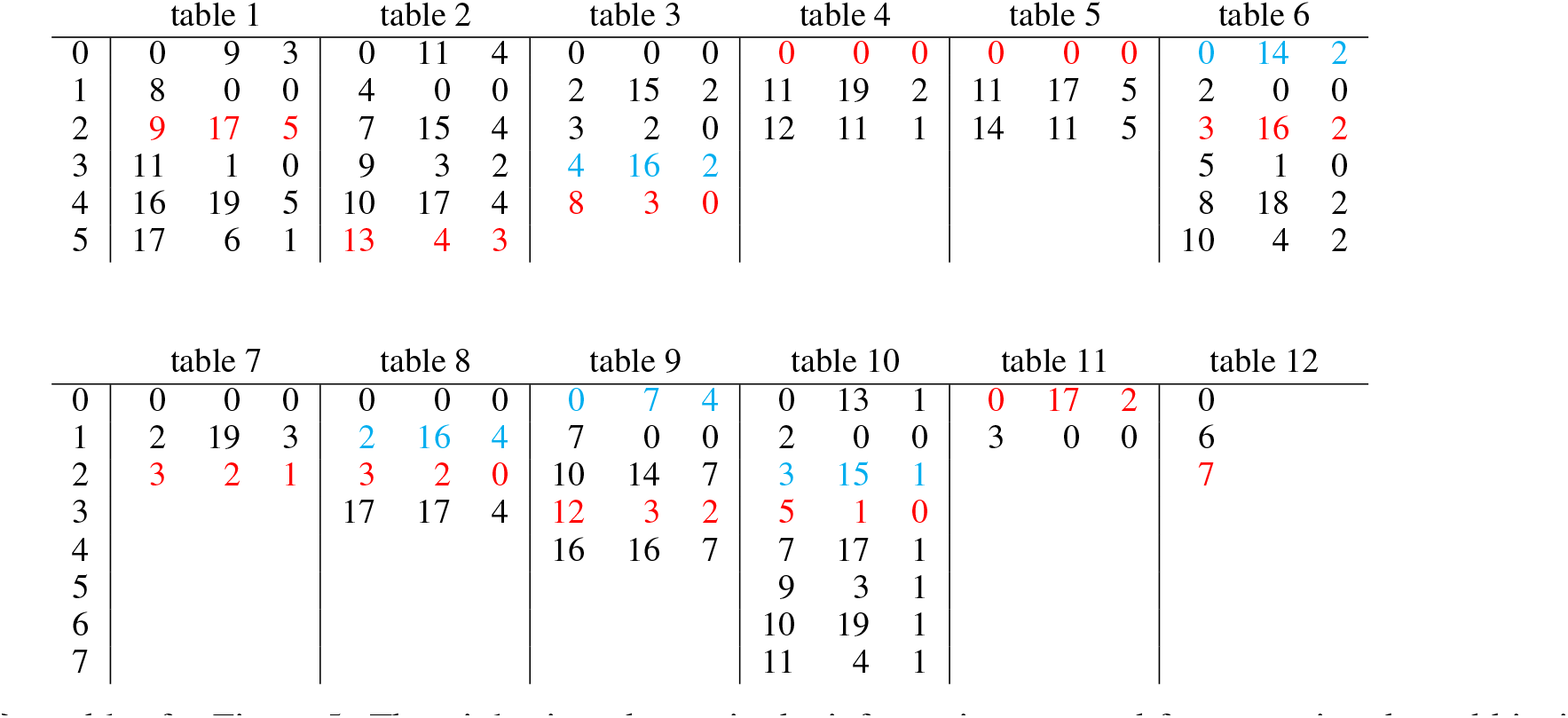
Our tables for Figure 5. The triples in red contain the information we need for extracting the red bits in Figure 5. If the triple of the *t*th column brings us to an entry at row *x* in the *t* + 1st column prior to a red line, we color the *x*th row in blue.

Looking at Figure 6, it is not hard to devise a variation of fractional cascading that limits the number of triples we scroll down in each table to 1, while increasing the total size of the tables by at most a factor of 2, as shown in Figure 7. For every other triple (*x, y, z*) in a table, we ensure there is triple (*x*′, *x, z*′) in the preceding table.

**Figure 7:**
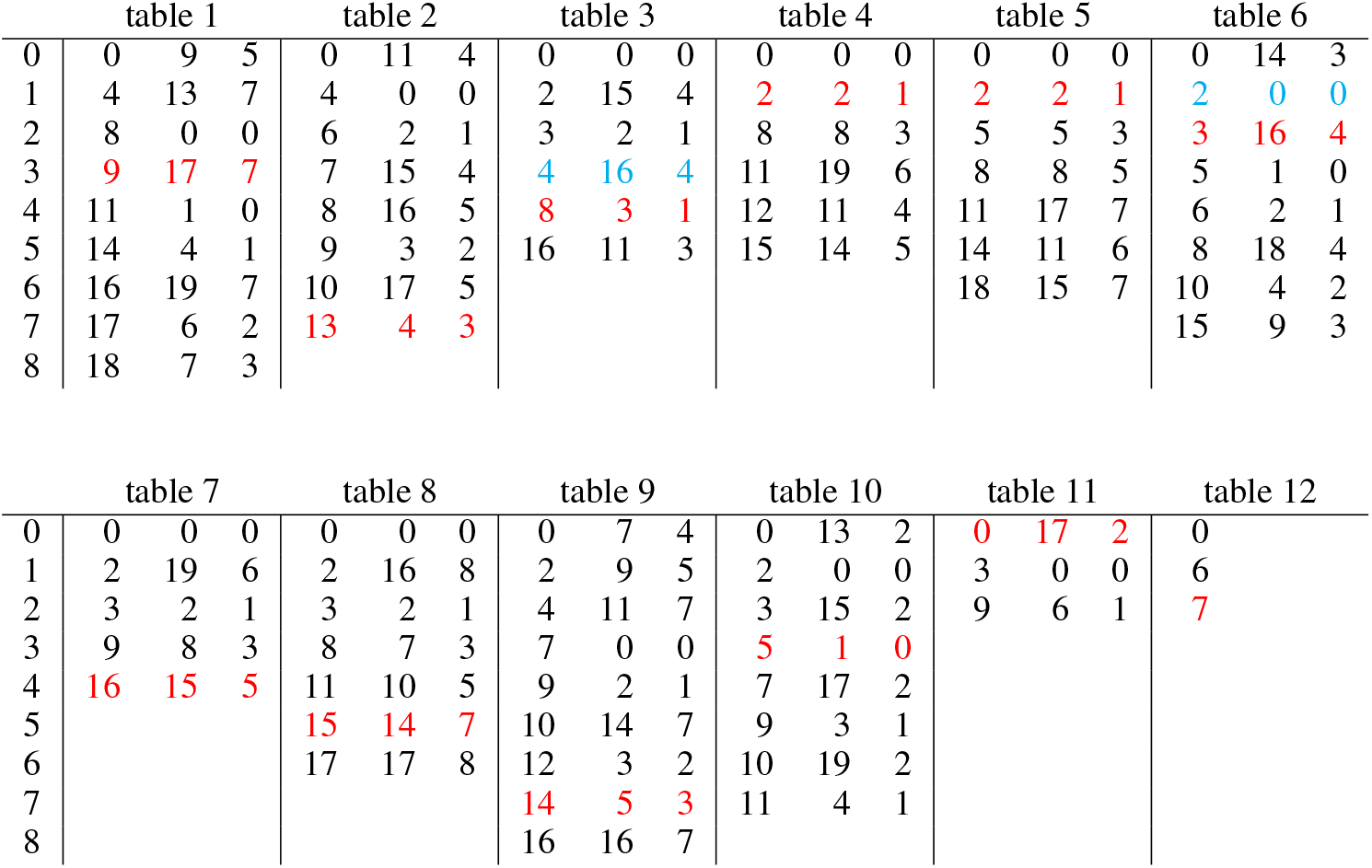
The tables in Figure 6 after we have applied our variation of fractional cascading. The triples in red again contain the information we need; those in blue indicate that the table lookup sends us to the preceding entry we want, so we must scan down only for one entry.

## Footnotes

1 The minimum LCP value may not be unique. For our purposes, we will assume we take the position of the first minimum LCP value.

